# Shear stress inhibits adipocyte differentiation via downregulating lncRNA MALAT1

**DOI:** 10.1101/2025.01.20.633971

**Authors:** Justin Caron, Madison Marino, Bo Wang, Shue Wang

## Abstract

Adipocyte differentiation plays an important role in bone remodeling due to secretory factors that can directly modulate osteoblast and osteoclast, thus affecting overall bone mass and skeletal integrity. Excessive adipocyte differentiation within bone marrow microenvironment can lead to decreased bone mass, eventually causing osteoporosis. Mechanical microenvironment of bone marrow, including fluid shear, maintains the balance of adipocyte and osteoblast differentiation during bone remodeling. However, how mechanical cues interact with long noncoding RNA (lncRNA) and regulating adipocyte differentiation remains unexplored. In this study, we investigated the mechanosensitive role of lncRNA MALAT1 during mesenchymal stem cells (MSCs) adipocyte differentiation. By applying physiologically relevant shear stress, MSCs experienced morphological changes and adipocyte differentiation differences. Shear stress inhibits MSCs adipocyte differentiation with reduced oil-red-o-stained lipid droplets. Silencing MALATs results in reduced adipocyte differentiation. By leveraging a novel gapmer double stranded locked nuclei acid (dsLNA) nanobiosensor, we showed that shear stress inhibits MALAT1 expression with significantly reduced fluorescence intensity. Our findings suggest that shear stress influences adipocyte differentiation primarily through the downregulation of MALAT1, highlighting a significant interplay between biophysical cues and lncRNAs. This interaction is crucial for understanding the complexities of bone remodeling and the potential therapeutic targeting of lncRNAs to treat bone-related disorders.

## Introduction

Mesenchymal stem cells (MSCs) are multipotent, self-renewing stem cells that are able to differentiate into numerous cell types, including osteoblasts, adipocytes, chondrocytes, hepatocytes, cardiomyocytes, and nerve cells with target specific biochemical factors ^1, 2^. Increased evidence has shown MSCs have been the favorite cell sources for the application of tissue engineering and regenerative medicine due to their remarkable properties. Various studies have shown that MSCs have been applied in repairing bone ^3, 4^, cartilage ^5^, adipose ^6^, nerve ^7^, and other organs ^8, 9^. However, the success of MSCs-based cell therapy and tissue engineering is highly dependent on MSCs fate determination or commitment. The uncontrolled MSCs differentiation can lead to unwanted differentiation of undesired phenotype, which limits their applications for tissue repair or regeneration. Thus, how to precisely control MSCs differentiation into desired lineage or phenotype is critical for MSCs-based therapy. Over the last few decades, unremitting efforts have been devoted to understanding biochemical signals that regulate MSCs commitment. A number of critical signaling pathways are identified and involved in regulating MSCs lineage commitment, including bone morphogenetic protein (BMP) signaling, Wnt signaling, and Notch signaling ^10–12^. Recently, accumulating data suggested that long non-coding RNAs (lncRNA) emerge as novel regulators of numerous biological processes, including embryonic development, cancer progression, and stem cell fate determination. lncRNAs are transcripts longer than 200 nucleotides that are not translated into proteins but play important regulatory roles in transcriptional and post-transcriptional regulation. Recent studies have shown that several lncRNAs are involved in the regulation of stem cell fate commitment ^13^. For example, it is reported that lncRNA-LULC active smooth muscle differentiation of adipose-derived MSCs by upregulation of BMP9 expression; lncRNA MALAT1 ^14^, MSC-AS1 ^15^, lncRNA-OG, H19 ^16^, NEAT1 ^17^ promote osteogenic differentiation either through miRNA-related regulation or chromatin remodeling ^18^. lncRNA HOTAIR ^19, 20^, GAS5 ^21^, and ANCR ^22^ were reported to negative regulate adipocyte differentiation. Over the past decade, studies have extensively examined how mechanical cues like shear, stiffness, and topography affect MSCs fate determination ^23, 24^. Fluid shear stress (SS), resulting from exposure to vascular or interstitial fluid flow, is a key biophysical factor that influences mesenchymal stem cell (MSC) fate. However, it has been studied less compared to other mechanical cues, despite its significant impact on MSC differentiation and behavior. Our recent study showed physiological relevant shear stress (3 - 7 dynes/cm^2^) enhance MSCs osteogenic differentiation and partially through Notch activation ^25^. However, the effects of shear stress on adipocyte differentiation are obscure and have controversial results. In a study by Adeniran-Catlett et al., the authors demonstrated that shear stress (15 dyne/cm^2^, 1.5 Pa) enhance adipogenesis with increased accumulation of lipids ^26^. However, in several other studies, it has been reported shear stress in the range of (6 – 10 dyne/cm^2^) suppressed MSCs adipogenesis as indicated by reduced lipid droplet accumulation ^27^. Interestingly, significantly lower shear stress (0.28 dyne/cm^2^) demonstrated a marked increase in lipid accumulation ^28^. Another study showed in obese patients, the wall shear stress of the vein is lower (lower than 0.25 Pa) compared to control group (0.3-0.6 Pa) ^29^. Thus, it is intriguing to study the effect of shear at the range of 0.3 – 0.6 Pa on adipogenesis of MSCs, which mimicking the normal physiological relevant shear. Inspired by native physiological microenvironment, recent studies suggest that shear stress can act in synergism with other biochemical cues to augment the differentiation efficiency of MSCs. This combination effectively replicates *in vitro* conditions of MSCs. Here, we investigated shear stress in combination of biochemical factor, lncRNA MALAT1 in regulating MSCs adipocyte differentiation. We first investigated and compared the effects of fluid shear stress on MSCs proliferation and adipocyte differentiation. The phenotypic behaviors, including morphology, proliferation, and differentiation were characterized and compared. We further studied the role of lncRNA MALAT1 in regulating adipocyte differentiation. Finally, we examined the mechanosensitive role of MALAT1 using a novel nanobiosensor. Our experimental results provide convincing evidence supporting that physiologically relevant shear stress inhibits adipocyte differentiation of MSCs with significantly reduced accumulation of lipid droplets. Silencing MALATs results in reduced adipocyte differentiation. Our study suggest that shear stress inhibits MSCs adipocyte differentiation through the downregulation of lncRNA MALAT1, as further validated by quantitative RT-PCR analysis. This study will add new information on mechanosensitive role of MALAT1 in regulating MSCs lineage determination. These results illuminate the critical roles of mechanical environments and genetic regulators in skeletal health and disease.

## Materials and methods

### Cell culture and reagents

Human bone marrow-derived MSCs were acquired from Lonza, which were originally isolated from normal adult human bone marrow withdrawn from bilateral punctures of the posterior iliac crests of volunteers. MSCs were maintained in basal medium MSCBM (Lonza) and supplemented with L-glutamine, GA-1000, and cell growth factor (Lonza). Cells were cultured in tissue culture dishes in a humidified incubator at 37 °C with 5% CO_2_ with medium change every three days. Cells were passaged using 0.25% EDTA-Trypsin (Invitrogen) and passage 2-7 were used for the experiments. For adipocyte differentiation, MSCs were seeded at a concentration of 400 cells/mm^2^ with a volume of 500 *μ*L basal medium in 24 well-plates. Cells were maintained in basal medium until they reached 70-80% confluency. Then the basal medium was replaced with adipocyte differentiation medium and refreshed every three days. The adipocyte differentiation was evaluated after 5, 7, and 10 days of induction. **Figure 1A** showed the experimental design process.

**Figure 1.**
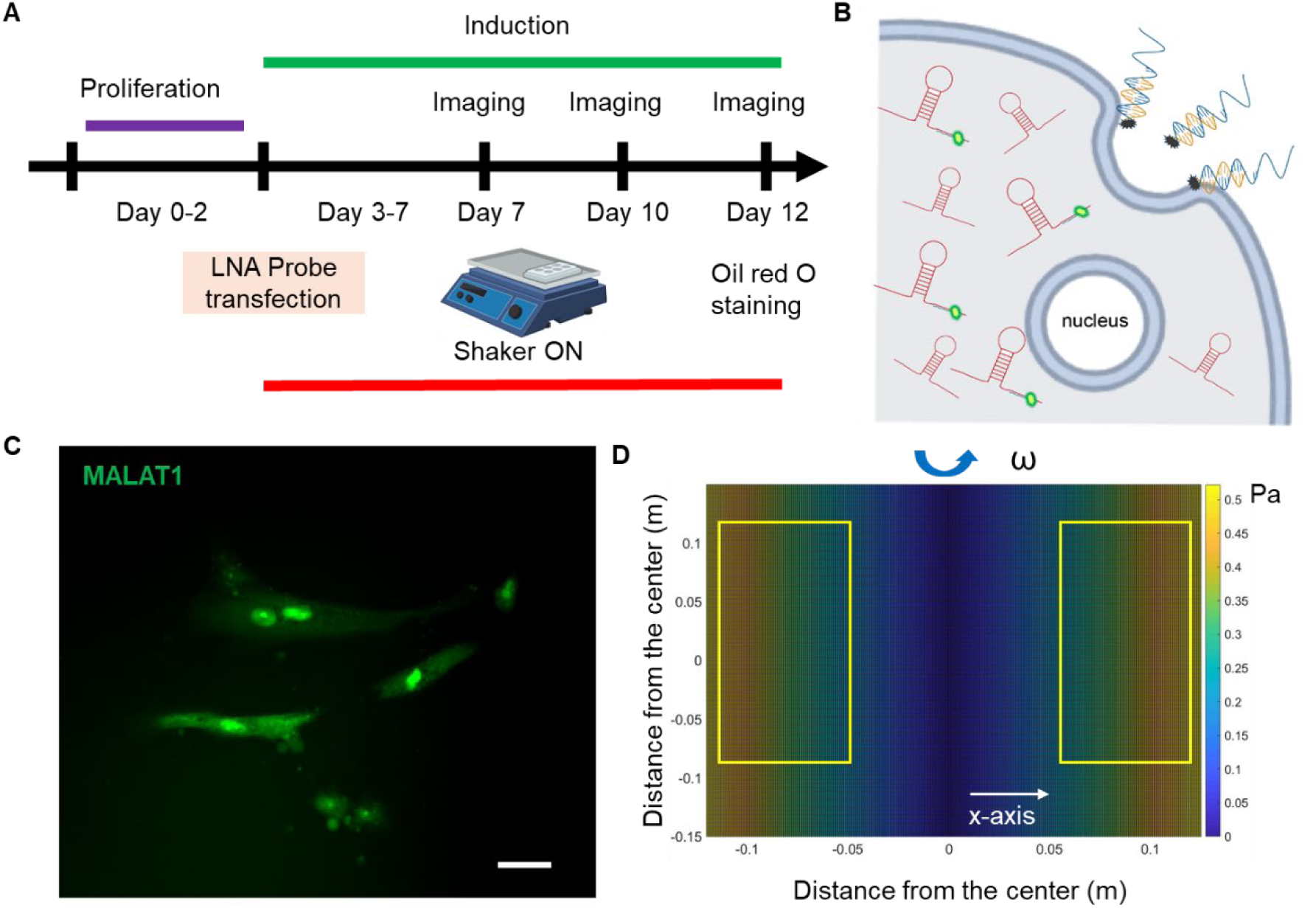
Overview of experimental design. **(A)** Experimental design. **(B)** Schematic illustration of endocytic uptake of LNA nanobiosensor by MSCs for lncRNA detection. Once internalized, the LNA detecting probe is displaced from the dsLNA complex and binds to the target lncRNA, allowing the fluorophore to fluorescence. **(C)** Representative fluorescence image of lncRNA MALAT1 expression in MSCs. Scale bar: 50 µm. **(D)** Simulated distribution of orbital shear stress. Yellow labeled rectangles indicate the location of well-plates. The estimated shear stress was in the range of 0.3 - 0.6 Pa.

### Preparation of double-stranded LNA probe

To prepare the LNA/DNA nanobiosensor, the LNA detecting probe and quencher probe were initially prepared in 1x Tris EDTA buffer (pH = 8.0) at a concentration of 100 nM. The LNA probe and quencher were mixed at the ratio of 1:2 and incubated at 95°C in a dry water bath for 5 min and cooled down to room temperature over the course of 2h. Once cooled down, the prepared LNA probe and quencher mixer can be stored in a refrigerator for up to 7 days. For lncRNA detection, the prepared double-stranded LNA/DNA probe was then transfected into MSCs using Lipofectamine 2000 following manufacturers’ instructions. lncRNA gene expression can thus be evaluated by measuring the fluorescence intensity of MSCs transfected with LNA/DNA probes.

### Simulation of orbital shear stress

To apply the simulated shear, a low-speed orbital shaker (Corning LSE, 6780-FP, orbit) was used. Once MSCs reach 90% confluency, cells were induced using adipocyte induction medium. One plate of cells was then placed on the orbital shaker at the speed of 10 RPM continuously for a total of 10 days. The orbital shaker was placed inside the cell culture incubator to maintain the cell culture environment. The orbital shear stress was calculated using the following equation: 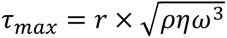, where *τ_max_* is near-maximal shear stress, *r* is the orbital radius of rotation, *ρ* is the density of cell culture medium, *η* is the dynamic viscosity of the medium, *ω* is the angular velocity and *ω=2πf*, where *f* is the frequency of rotation (revolution per second).

### Silencing MALAT1

To silence lncRNA MALAT1 expression, MSCs were seeded in 12-well plates and transfected with MALAT1 antisense probe (Qiagen) at a concentration of 50 nM (final concentration) using Lipofectamine 2000 (Invitrogen) transfection reagent following manufacturer’s instructions. A negative control antisense probe was used as a negative control. After 24 hours of transfection, the transfection medium was replaced with a fresh basal culture medium. To investigate the effects of silencing MALAT1 on adipocyte differentiation, induction was initiated after 24 hours of MALAT1 silencing.

### Reverse Transcription and RT-PCR

To quantify the silencing efficiency of MALAT1 in MSCs, the RT-PCR assay was performed. Initially, cells were seeded in 6-well plates with a concentration of 4 x 10 ^5^ cells/well. After siRNA silencing for 48 hours, total RNAs were isolated and reverse-transcribed into cDNA using SuperScript VILO cDNA Syntheis Kit (ThermoFisher, Cat #: 11754050). cDNA samples were then amplified by qPCR. PCR reaction solution was assembly as below: 0.5 µL Taqman Gene Expression Assay (10x) for MALAT1 (Assay ID: Hs00273907_s1, Cat #: 4331182), 5 µL of TaqMan Fast Advanced Master Mix (Cat #:4444556), and 1 µL of cDNA. The total volume is 10 uL. The quantitative PCR was performed on a BioRad Real Time PCR system, and data were collected and analyzed. All samples were prepared and tested in triplicate. The relative expression levels of lncRNAs were determined by equation 2*^−ΔΔ^*^Ct^.

### F-actin staining

The MSCs were washed 3 times with 1x PBS, 4 minutes each time. Then the cells were fixed with 4% paraformaldehyde solution (PFA) for 10 min before being permeabilized and blocked with the Perm/Block solution PBST (PBS + 0.5% Triton + 1% BSA) for 1 h. After washing 3 times with 1x PBS, cells were incubated with phalloidin (1:30) and Hoechst 33342 (1:2000) for 30 minutes at room temperature. The cells were then washed 3 times with 1x PBS before fluorescence imaging and analysis.

### Live/dead viability staining

The viability of MSCs following exposure to orbital shear was assessed using a live/dead viability assay (ThermoFisher). Cells were stained with propidium iodide (PI, 10 *μ*g/ml), a fluorescent agent that binds to DNA by intercalating between base pairs. Hoechst 33342 was used to stain cell nuclei at a concentration of 20 *μ*M for 30 minutes. After staining, cells were washed three times with 1x PBS to remove excess dye. Imaging was performed using Texas Red (535/617 nm) and DAPI (360/460 nm) filters on the ZOE imaging station.

### Oil Red O staining

To assess adipocyte differentiation of MSCs, intracellular lipid droplets, which are markers of adipocyte formation during differentiation, were stained using Oil Red O. The working solution were prepared by diluting Oil Red O stock solution 3:2 using deionized H_2_O (diH_2_O) and filtered using filter paper. The staining process were performed based on the manufacturer’s instructions. Briefly, after 5, 7, and 10 days of adipocyte differentiation, MSCs were fixed using 4% PFA for 15 minutes and then washed 3 times with diH_2_O, followed by incubation of Oil Red O working solution for 15 minutes at room temperature. MSCs were then washed 5 times with diH_2_O and imaged using bright fields. Positive Oil Red O staining appears as red-stained lipid droplets within the cells, indicating successful adipocyte differentiation. The number of Oil Red- O positive lipid droplets were counted. Droplets with faint colors were ignored.

### Imaging and statistical analysis

Images were obtained using Echo Revolution fluorescent microscope with an integrated digital camera (5MP CMOS Color for bright field, 5 MP sCMOS Mono for fluorescence imaging). Actin staining images were acquired using a ZOE Fluorescent Cell Imaging Microscope (Bio-Rad). To ensure consistency, all images were captured under identical settings, including exposure time and gain. Image analysis and data collection were performed using NIH ImageJ software. To measure lncRNA MALAT1 expression, the mean fluorescence intensity of each cell was measured, and background noise subtracted. Cells were quantified within the same field of view, with a minimum of five images analyzed for each condition. The experiments were conducted at least three times, with over 100 cells quantified per group. Results were analyzed using independent, two-tailed Student t-test in Excel (Microsoft). P < 0.05 was considered statistically significant.

## Results

### Design of dsLNA nanobiosensor for lncRNA detection

To investigate the role lncRNA MALAT in adipocyte differentiation of MSCs that were exposed to shear stress, we utilized a double-stranded LNA (dsLNA) nanobiosensor for lncRNA MALAT1 expression analysis. The design and characterization of this dsLNA nanobiosensor was reported in our previous study ^30^. Briefly, this dsLNA nanobiosensor is a complex of a detection LNA probe and a quencher probe, with the length of 30- and 15- oligonucleotide sequence, respectively. The detection probe is a single-stranded LNA/DNA, with LNA monomers at the two ends of the oligonucleotide sequence that was designed to be complementary of partial MALAT1 sequence, **Table. S1**. The sequence was selected and validated for its specificity using RNA fold and BLAST ^30^. The quencher probe is a 15-base pair single-stranded LNA/DNA, with a quenching dye at the 3’ end of the sequence. The detection probe will bind to the quencher probe to form stable dsLNA complex, initiating the quenching. Once the LNA complex is transfected into the cells, in the presence of the lncRNA target sequence, the LNA detection probe is thermodynamically displaced from the quencher, allowing the fluorophore to fluorescence, thus detecting lncRNA expression at the single cell level, **Figure 1B-1C**. This displacement is due to a larger difference in free binding energy between LNA detection probe to target lncRNA versus LNA probe to quencher.

### Simulation of orbital shear stress and analysis

To simulate physiologically relevant shear stress and access its effects on adipocyte differentiation, an orbital shaker was utilized to simulate shear stress. We estimated the generated shear using Stokes’ second problem, which describes a plate oscillating along a single axis in the plan of the plate, with liquid above it. Although the shaker does not generate uniform laminar flow on cells, most cells were subjected to near-maximum shear, calculated as: 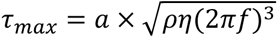, where *a* is the orbital radius of rotation, *ƞ* and *ρ* are viscosity and density of the culture medium, respectively. We further simulated the distribution of shear stress across the shaker platform using MATLAB to identify the slightly difference of shear stress across the entire platform. As the shaker oscillates along the y-axis, the generated shear stress along this axis remains consistent. The orbital shear was simulated when the shaker was setting at 30 RPM, with maximum shear stress of approximately 0.7 Pascal (7.1 dyne/cm^2^) at the edge of the shaker. The well plates were placed on the yellowed labeled region on top of the shaker, **Figure 1D**. The applied shear stress in different wells ranges from 3 to 7 dyne/cm^2^, which is consistent with the value reported in previous studies ^31, 32^. This range provides a realistic simulation of the mechanical forces acting on cells in the biological setting.

### Shear stress modulates MSCs morphology and actin organization

The effects of shear stress on MSCs phenotypic behaviors were first evaluated and compared, including cell viability, cell proliferation, cell aspect ratio, and cell perimeter. The cell viability was assessed using PI/Hoechest 33342 staining assay. **Figure S1-S2** show the bright field and fluorescent images of MSCs after 5 days and 10 days of culture w/o shear stress, respectively. It is evident that shear stress did not have effects on cell viability, with no dead cells observed in red channel. The effects of shear stress on cell morphology were further assessed by measuring cell aspect ratio and cell perimeter, respectively. **Figure S3A** showed actin cytoskeleton (stained in red), the nuclei (stained in blue), the merged images (actin + nuclei). It is shown that the cells under shear stress (bottom row) appear to have more elongated, aligned actin fibers compared to the control, suggesting that shear stress may influence actin cytoskeleton organization. Quantitative analysis of aspect ratio and perimeter measurements did not show significant differences between the control and shear-exposed cells, suggesting shear stress did not affect the overall cell shape or size in terms of perimeter or aspect ratio. This result suggests that shear stress might influence intracellular organization without drastically changing the overall size or shape of the cells. We next compared cell morphology changes when cells were cultured in adipocyte induction medium. Interestingly, shear stress affects cell aspect ratio and perimeter significantly, **Figure 2**. **Figure 2A** showed representative bright field and fluorescence images of MSCs under static and shear condition, respectively.

**Figure 2.**
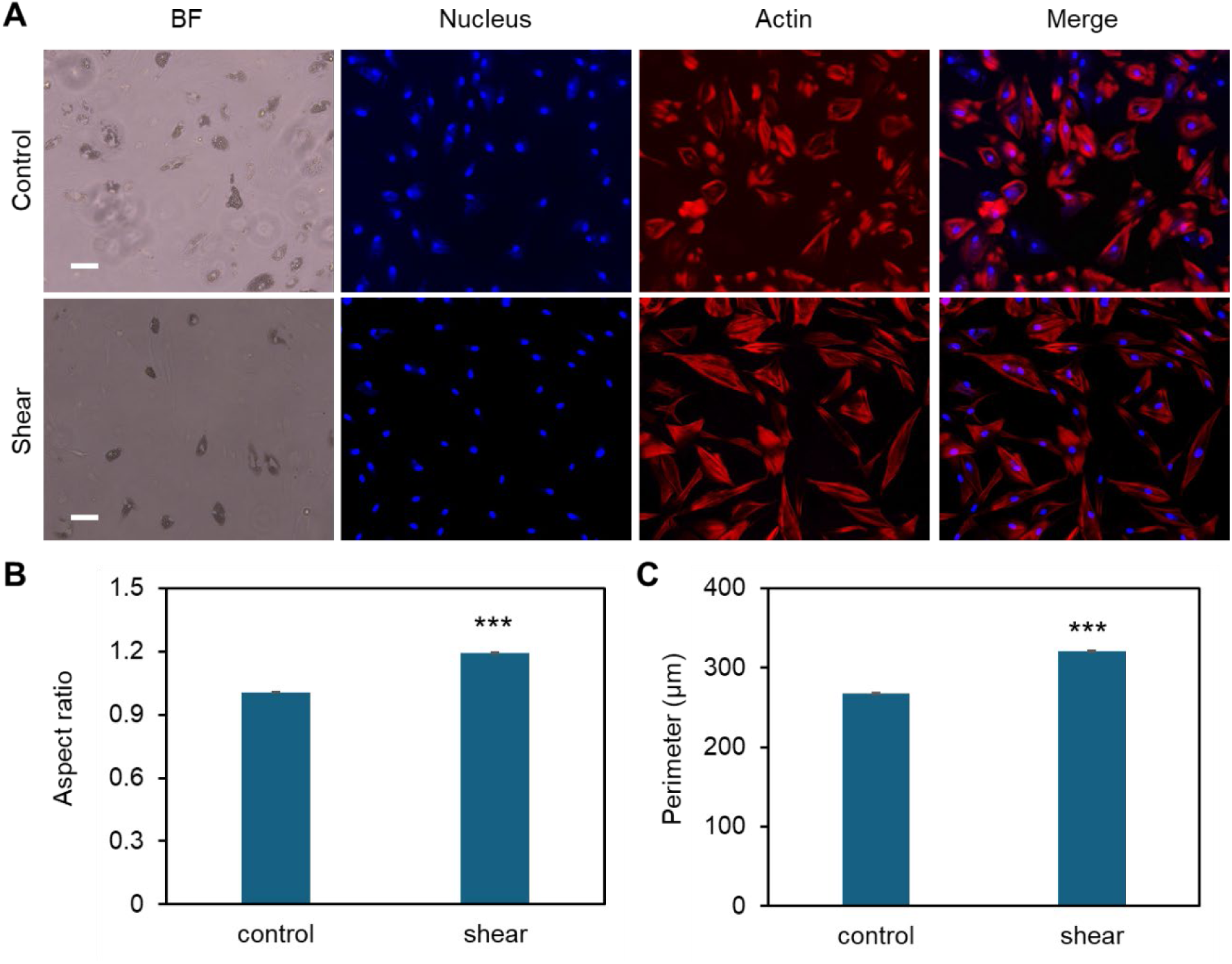
Effects of low fluid shear on MSCs morphology change during adipocyte differentiation. **(A)** Representative bright field and fluorescence images of MSCs under static condition (control) and exposed to shear stress (shear). MSCs were exposed to orbital shear continuously for 5 days. Samples were stained with F-actin (red; by phalloidin), and nuclei (blue; by Hoechst 33342), respectively. Scale bar: 100 μm. **(B)** Comparison of aspect ratio of MSCs after 10 days of exposure to low fluid shear. **(C)** Comparison of observed cell perimeter after 10 days of exposure to shear. Data represents over 200 cells in each group and are expressed as mean± s.e.m. (n = 5, ***, p < 0.001, **, p < 0.01, *, p < 0.05).

The cell aspect ratio and perimeter of cells that were exposed to shear stress increased significantly, **Figure 2B-2C**. It is noted that actin cytoskeleton is under remodeling when MSCs cultured in adipocyte differentiation medium, **Figure S4**. In the control group, the actin filaments were less organized compared to the cells exposed to shear. However, the actin filaments were still well organized even after 5 days of induction, indicating the strong shear effects on actin organization.

### Shear stress inhibits adipocyte differentiation

Preciously, we have reported that low fluid shear stress enhances MSCs osteogenic differentiation with enhanced ALP enzyme activity ^25^, however, it is obscure how shear affects adipocyte differentiation. Thus, the effects of shear stress on MSCs adipocyte differentiation were further elucidated and compared. Briefly, cells were initially seeded in two well plates and cultured in basal medium under static condition. Once the cells reached 70–80% confluency, basal medium was replaced by induction medium for both plates. Meanwhile, one well plate was placed on top of the orbital shaker, while the other plate was placed in static condition without exposure to shear. After 5, 7, and 10 days of induction and shaking, adipocyte differentiation was evaluated and compared by measuring oil-red-o-stained cells, a biochemical marker for lipid accumulation, a crucial step for adipocyte formation. **Figure 3A** showed the representative MSCs images after 5, 7 and 10 days of adipocyte induction in control and shear condition, respectively. We further quantified the differentiated cells by counting the oil-red-o-stained adipocyte lipid droplets in each field of view. **Figure 3B** showed comparison of differentiated cells at different time points, with and without shear stress. It is evident that for the cells without shear stress, the number of differentiated cells increases as the days increase.

**Figure 3.**
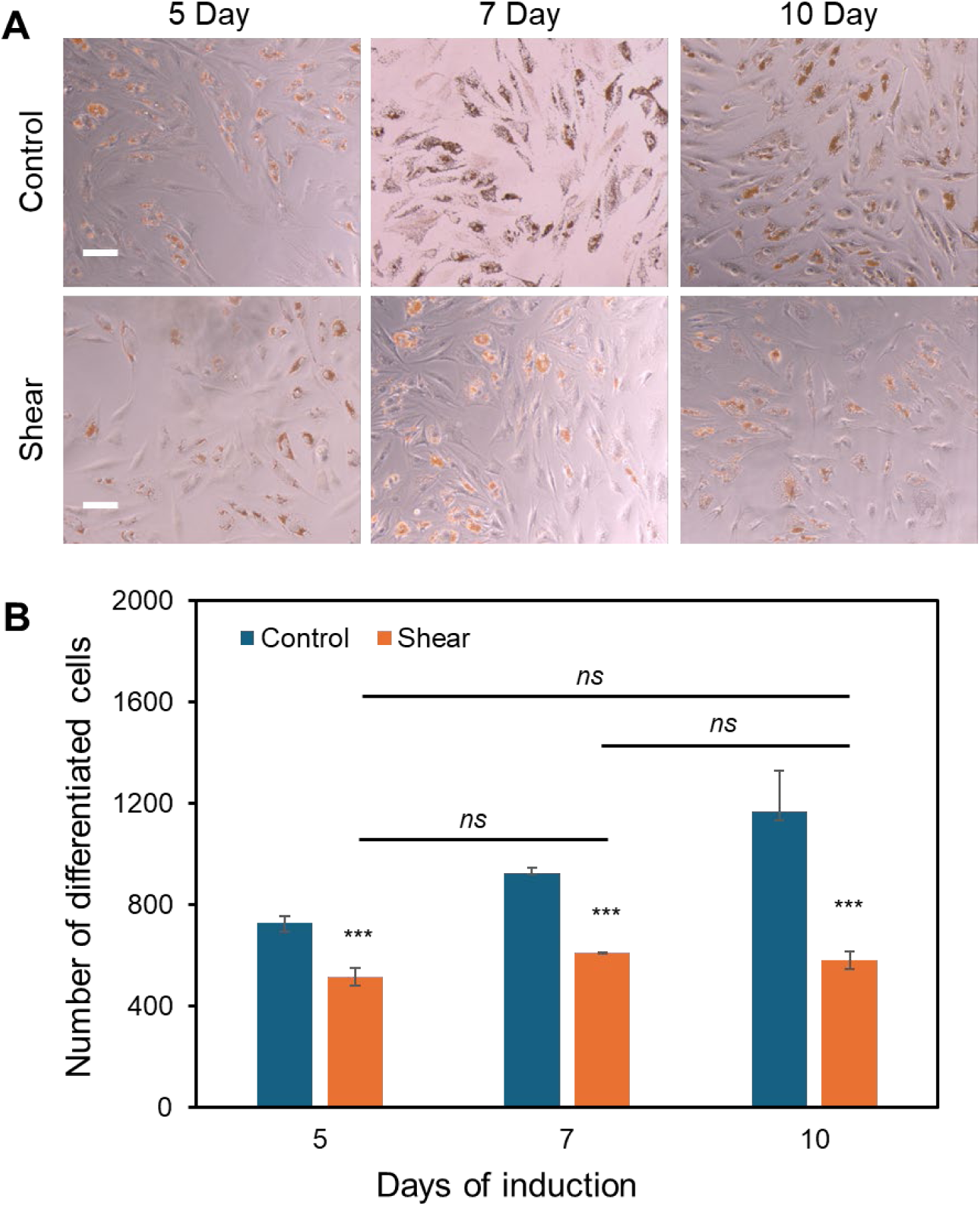
Low fluid shear inhibits adipocyte differentiation. **(A)** Representative images of adipocyte differentiation after 5, 7, and 10 days with and without shear. **(B)** Quantitative results of adipocyte comparison. The differentiated adipocytes were quantified and compared by measuring the number of lipid droplets at each field of view. Data represents over 500 cells in each group and are expressed as mean± s.e.m. (n = 5, ***, p < 0.001, **, p < 0.01, *, p < 0.05).

However, for the cells exposed to shear stress, the number of differentiated cells was significantly decreased, and the number was not increased after 7 and 10 days of induction. Together with the previous observation, the results indicate that shear stress modulates cell morphology, proliferation, and inhibits adipocyte differentiation.

### MALAT1 knockdown inhibits adipocyte differentiation of MSCs

It had been reported that lncRNA MALAT1 enhances osteogenic differentiation of bone-derived and adipose-derived MSCs through either sponging to miRNAs by enhancing downstream genes expression, including Runx2, ALP, and BMP ^33^. Our group also showed earlier MALAT1 enhance osteogenic differentiation with increased ALP activity and calcium deposition.

However, the role of MALAT1 in adipocyte differentiation was not well explored, especially the crosstalk of MALAT1 and biophysical cues. Thus, we investigated the role of MALAT1 in adipocyte differentiation and the effects of shear stress. We first silenced MALAT1 using siRNA knockdown. A negative control siRNA was used as a control for comparison. Briefly, cells were seeded in well plates and silenced using control siRNA and MALAT1 siRNA, following manufacture’s instructions. The silencing efficiency was compared, **Figure S5**. To evaluate the effects of shear stress on MALAT1 silenced cells, the cells were placed on the shaker for the duration of adipogenic induction. **Figure 4A** showed representative images of differentiated MSCs in different conditions after 5 and 10 days of induction. The results showed silencing MALAT1 inhibits adipocyte differentiation with reduced lipid droplets and nodules after 5- and 10- days induction. Without applying shear, MALAT1 siRNA knockdown reduced the number of differentiated cells by 62% and 65% after 5 and 10 days of induction, respectively **(Figure 4B)**. For the cells exposed to shear stress, MALAT1 knockdown reduced adipocyte differentiation by 54% and 65% after 5- and 10- days of induction, respectively **(Figure 4B)**. It is noted that applying shear stress did not further inhibit adipocyte differentiation for the MALAT1 siRNA silenced groups. This result indicates that shear stress may downregulate the expression of MALAT1, leading to a reduction in lipid droplet accumulation within cells.

**Figure 4.**
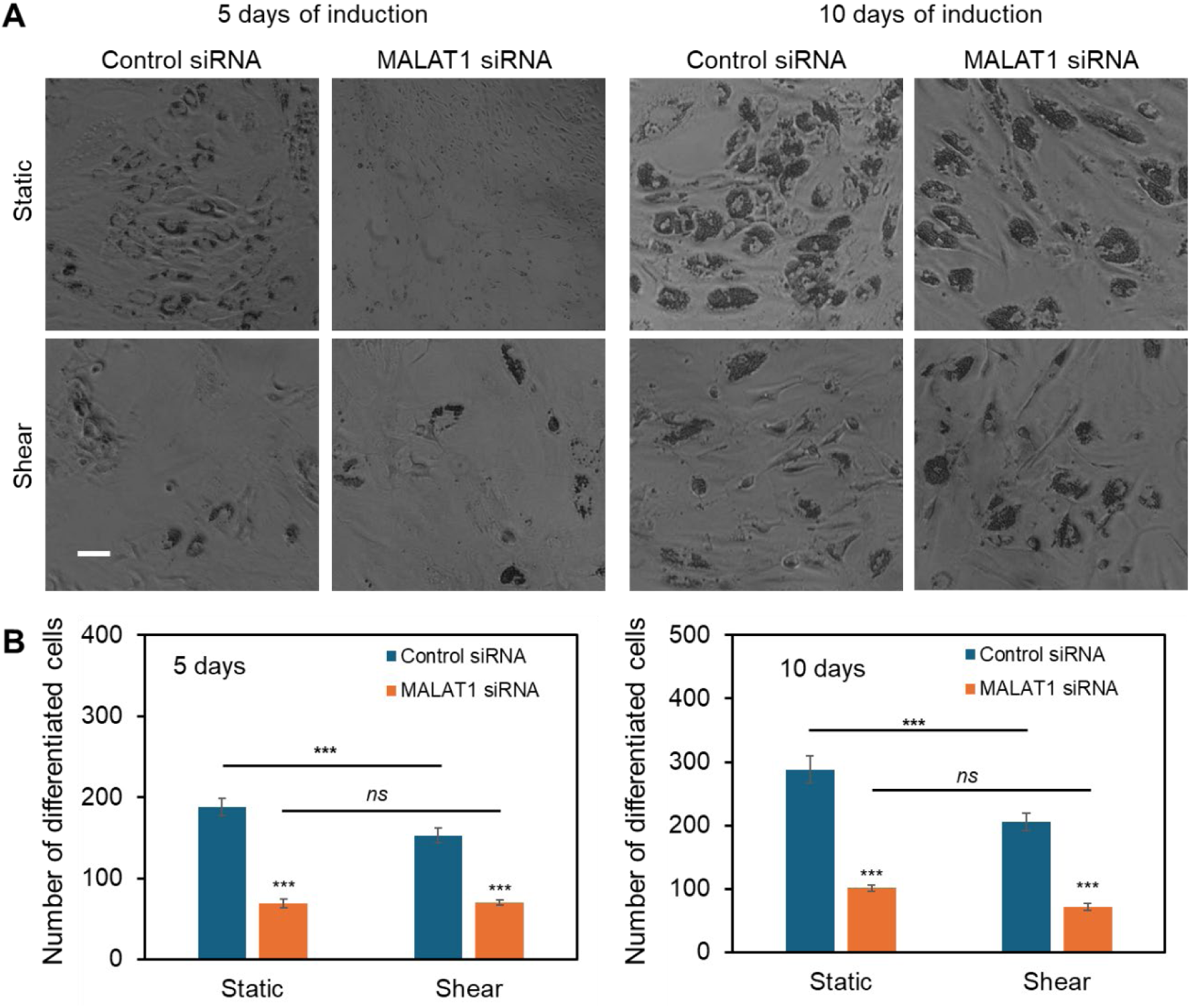
lncRNA MALAT1 knockdown inhibits adipocyte differentiation. **(A)** Representative images of differentiated MSCs in different conditions (static and shear) after 5 and 10 days of induction. Scale bar: 100 µm. **(B)** Comparison of the number of differentiated cells after MALAT1 silencing for both cells under static and shear conditions (n=3). Data are expressed as mean ± s.e.m. (n = 3). A two-tailed t-test was used to analyze differences between control and shear conditions. *, p < 0.05; **, p < 0.01; ***, p < 0.005.

### Shear stress modulates the expression of MALAT1

Previous studies indicate that lncRNAs are responsive to biophysical cues, indicating their mechanosensitive roles in regulating cell functions, including proliferation, migration, and differentiation. Wu et al. showed that lncRNA H19 participate in osteogenic differentiation by sponging miR-135 to regulate MSCs activities via FAK in a tension-induced way ^34^. A new lncRNA that is related to mechanical stress (lncRNA-MSR) would hijack miR-152 to control TMSB4 expression in cyclic tensile strain-induced regulation of chondrocytes and enhance cartilage degradation ^35^. Thus, to test whether MALAT1 is responsive to shear stress, we utilized an LNA/DNA gapmer nanobiosensor to detect MALAT1 expression in MSCs ^30^. **Figure 5A** showed the bright field and fluorescence images of MSCs under control and shear conditions, respectively. The results showed that shear stress inhibits MALAT1 expression with reduced fluorescence signal. We further quantified the inhibition effects by measuring the mean fluorescence intensity of each individual cells and compared, **Figure 5B**. For the cells exposed to shear stress, the expression of lncRNA MALAT1 was reduced by approximately 40% compared to the cells without shear. To validate shear stress inhibits MALAT1 expression, we performed RT-PCR to quantitatively evaluate MALAT1 expression in both control and shear-exposed cells, **Figure 5C**. The results showed that MALAT1 expression was downregulated when the cells exposed to shear stress, with approximately 30% decrease. Taken together, the results shown MALAT1 is responsive to mechanical shear stress, indicating its mechanosensitive role in stem cell fate determination.

**Figure 5.**
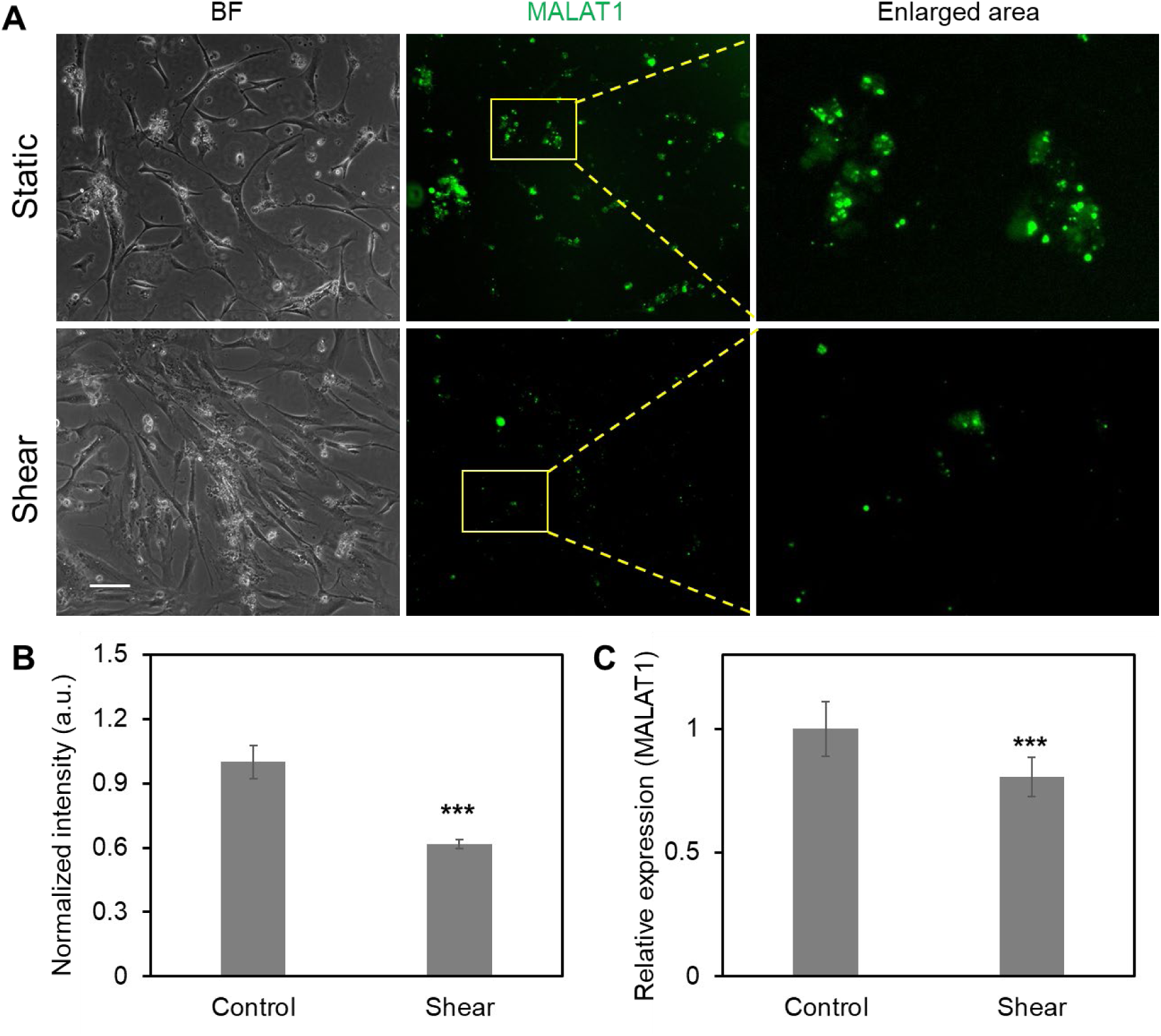
Shear stress modulates the expression of MALAT1. **(A)** Representative bright field and fluorescence images of MSCs in control and shear conditions, respectively. Green fluorescence signal shows MALAT1 expression. Scale bar: 100 µm. **(B)** Comparison of mean fluorescence intensity of MALAT1 expression (n=4). **(C)** Comparison of MALAT1 expression using RT-PCR assay. Experiments were performed at least three times. The relative expression levels of lncRNAs were determined by the equation 2^−ΔΔCt^. Data are expressed as mean ± s.e.m. (n = 3). A two-tailed t-test was used to analyze differences between control and shear conditions. *, p < 0.05; **, p < 0.01; ***, p < 0.005.

## Discussion

Biophysical and biochemical cues are key parameters that regulate cell behaviors, including proliferation, motility, and differentiation. Emerging evidence has shown that biophysical cues, including shear, stretch, and stiffness is pivotal in directing stem cell fate determination.

Although numerous studies have shown that stem cells are mechanosensitive and respond to mechanical stimuli, it is obscure which genome is directly responsive the stimulation. In this study, we first investigated the effects of shear stress on MSCs behavior, including cell proliferation and adipocyte differentiation. We next studied the role of lncRNA MALAT1 in MSCs differentiation and elucidated the correlation between shear stress and the expression of MALAT1 in MSCs. We discovered that MALAT1 expression was downregulated when cells were exposed to shear stress, indicating its mechanosensitive role. By utilizing a novel gapmer dsLNA nanobiosensor, we detected and compared MALAT1 expression in MSCs. This nanobiosensor allows for the monitoring of lncRNA expression in live cells at the single-cell level without requiring cell lysis or fixation. Previous studies have shown that this dsLNA nanobiosensor can track spatiotemporal RNA dynamics in cell migration ^36^, mice lung cancer ^37^, wounded corneal tissue repair ^38^, liver tissue ^39^, synthetic cells ^40, 41^, and vasculature formation ^42, 43^. Our group recently showed a dsLNA nanobiosensor in monitoring Dll4 dynamics during osteogenic differentiation ^25, 44^. It is noted that this nanobiosensor can be utilized to detect other types of RNA detection, i.e., miRNA and long non-coding RNA (lncRNA) ^30, 41, 45, 46^. Previous studies have demonstrated its capability in monitoring miRNA and mRNA dynamics during cell migration, 3D cancer invasion, and 3D osteogenic differentiation ^30, 46, 47^. This nanobiosensor is versatile, functioning effectively across various types and tissue environments.

Adipocyte differentiation plays a crucial role in bone remodeling because adipocytes within bone marrow secrete various factors that can directly modulate the activity of osteoblasts and osteoclast, thus affecting overall bone mass and skeletal integrity ^48^. Imbalanced adipocyte differentiation and osteoblast differentiation within bone marrow microenvironment could potentially lead to bone diseases. Excessive adipocyte differentiation can lead to decreased bone mass, contributing to conditions like osteoporosis, the most common bone remodeling disorder worldwide ^49, 50^. For example, increased marrow fat content has been observed in most bone loss conditions. In addition, bone marrow adipose tissue alterations have been associated with systemic conditions like anemia ^51^, glucose intolerance, and cancer ^52^. It was also reported that exposure to space with removed physical forces causes a 10- fold increase in adipocyte differentiation in animal models ^50^. The tightly controlled lineage commitment of MSCs plays a critical role in maintaining bone homeostasis, which is essential for proper function and balance of bone tissue. Therefore, understanding lineage commitment of MSCs to adipocyte is essential and could provide effective therapeutic regime for related bone diseases. In bone marrow microenvironment, bone cells sense mechanical load in response to fluid shear stress, initiating a signal for cellular excitation ^53^. The fluid flow is critical for mechanotransduction as well as enhancing convective solute transport within the bone ^54^. Thus, we investigated the effects of fluid shear affects MSCs adipocyte differentiation. Our results showed that in a simulated physiological shear environment, adipocyte differentiation is inhibited. Taking together with our previous finding that shear stress enhance osteogenic differentiation ^25^, these results indicate that bone marrow origin of MSCs has the potential to differentiate into both osteoblast and adipocyte and maintain a balance between them.

MALAT1, one of the few highly conserved nuclear long noncoding RNA, is abundantly expressed in normal tissues. MALAT1 has been shown to regulate gene expression at transcriptional and post-transcriptional levels, modulating chromatin organization, splicing, and mRNA stability. Initially, MALAT1 was indicated in tumorigenesis, metastasis, and cardiovascular diseases. Recently, emerging evidence showed MALAT1 plays a crucial role in bone remodeling and bone related diseases. Zhao et al. reported that MALAT1 deficiency promotes osteoporosis and bone metastasis of melanoma, thus they concluded that MALAT1 protects against osteoporosis and bone metastasis ^55^ . Several studies also showed low expression of MALAT1 is related to reduced osteoblast differentiation and cause bone mass loss ^56, 57^. Nevertheless, how MALAT1 responses to mechanical cues, i.e., shear stress, has not been explored. Based on literature studies and our previous findings, we hypothesize that MALAT1 is a mechanosensitive biomarker that regulates bone remodeling by controlling osteogenic and adipocyte differentiation. We further verified that MALAT1 expression was downregulated in MSCs that were exposed to shear stress. Without shear stress MSCs were induced to adipocyte differentiation, which is consistent with previous findings that reduced shear stress or altered shear stress promote adipocyte differentiation, which eventually causes reduced osteoblast differentiation, thus altering the bone homeostasis.

The differentiation of MSCs is a two-step process that include lineage commitment, which transitions MSCs into lineage-specific progenitors, and maturation, evolving progenitors into specific cell types. Several key regulators have been identified in regulating MSCs lineage commitment, including transforming growth factor-beta (TGF*β)/*bone morphogenic protein (BMP) signaling ^58^, wingless-type MMTV integration site (Wnt) signaling ^59^, Hedgehog (Hh) ^60^, and Notch signaling ^61, 62^. Although our study showed shear stress inhibits adipocyte differentiation through downregulating MALAT1, the downstream signaling pathway is not explored. Thus, future mechanistic studies will focus on elucidating the downstream signaling pathways. Specifically, MALAT1 may function through sponging miRNAs to regulate downstream gene expression, thus modulating adipocyte differentiation.

## Conclusion

In this study, we aimed to identify the mechanosensitive role of lncRNA MALAT1 during MSCs adipocyte differentiation. We first evaluated the physiological relevant shear stress on cell behaviors including morphological changes and differentiation capacity. Our results showed that cell viability and cell morphology were not affected by shear stress. However, once MSCs went through adipocyte differentiation, cells exposed shear showed elongated morphology with increased cell aspect ratio and perimeters. Interestingly, shear stress inhibits adipocyte differentiation with reduced lipid droplets and nodules. By disrupting the expression of lncRNA MALAT1, we further assessed the role of MALAT1 in regulating adipocyte differentiation. We showed silencing MALAT1 significantly mediated adipocyte differentiation. Thus, it is intriguing to study how shear stress modulates MALAT1 expression. By utilizing a novel gapmer dsLNA nanobiosensor, we monitored MALAT1 expression in MSCs and discovered that shear stress inhibits MALAT1 expression with significantly reduced fluorescence intensity. Quantitative RT-PCR analysis further validated this downregulation. Taken together, we conclude that shear stress inhibits MSCs adipocyte differentiation through downregulation of lncRNA MALAT1, indicating the crosstalk between biophysical cues and lncRNAs, and their important role in bone remodeling.

## Author contributions

S.W. and B.W. conceived the initial idea of the study. J.C. and M.M. performed the experiments.

S.W. and B.W. contributed to the experimental design and data analysis. J.C, M.M, B.W. and

S.W. contributed to the writing. S.W. finalized the manuscript with feedback from all authors.

## Acknowledgment

S.W. would like to acknowledge the support from the NSF CMMI program (CAREER Award: 2143151). J. Caron is supported by the Provost Graduate Fellowship.

## References

1. M. F. Pittenger, D. E. Discher, B. M. Péault, D. G. Phinney, J. M. Hare and A. I. Caplan, NPJ Regenerative medicine, 2019, 4, 1-15.

2. Y. K. Wang and C. S. Chen, Journal of cellular and molecular medicine, 2013, 17, 823–832.

3. H. Lin, J. Sohn, H. Shen, M. T. Langhans and R. S. Tuan, Biomaterials 2019, 203, 96–110.

4. Y. Zhang, Y. Xing, L. Jia, Y. Ji, B. Zhao, Y. Wen and X. Xu, Stem cells and development, 2018, 27, 1634–1645.

5. N. Fahy, M. Alini and M. J. Stoddart, Journal of Orthopaedic Research®, 2018, 36, 52–63.

6. S.-N. Li and J.-F. Wu, Stem cell research & therapy, 2020, 11, 41.

7. M. Fernandes, S. G. Valente, R. G. Sabongi, J. B. G. Dos Santos, V. M. Leite, H. Ulrich, A. A. Nery and M. J. da Silva Fernandes, Neural regeneration research, 2018, 13, 100.

8. Y. Han, X. Li, Y. Zhang, Y. Han, F. Chang and J. Ding, Cells, 2019, 8, 886.

9. J. K. Leach and J. Whitehead, ACS biomaterials science & engineering, 2017, 4, 1115–1127.

10. F. Guilak, D. M. Cohen, B. T. Estes, J. M. Gimble, W. Liedtke and C. S. Chen, Cell stem cell, 2009, 5, 17–26.

11. Y.-K. Wang, X. Yu, D. M. Cohen, M. A. Wozniak, M. T. Yang, L. Gao, J. Eyckmans and C. S. Chen, Stem cells and development, 2012, 21, 1176–1186.

12. S.-J. Heo, T. P. Driscoll, S. D. Thorpe, N. L. Nerurkar, B. M. Baker, M. T. Yang, C. S. Chen, D. A. Lee and R. L. Mauck, Elife, 2016, **5**, e18207.

13. C. Lanzillotti, M. De Mattei, C. Mazziotta, F. Taraballi, J. C. Rotondo, M. Tognon and F. Martini, Frontiers in Cell and Developmental Biology, 2021, 9, 742.

14. J. Yi, D. Liu and J. Xiao, Cell and tissue research, 2019, 376, 113–121.

15. N. Zhang, X. Hu, S. He, W. Ding, F. Wang, Y. Zhao and Z. Huang, Biochemical and biophysical research communications, 2019, 519, 790–796.

16. Z. Zhou, M. S. Hossain and D. Liu, Stem Cell Research & Therapy, 2021, 12, 1–9.

17. Y. Zhang, B. Chen, D. Li, X. Zhou and Z. Chen, Pathology-Research and Practice, 2019, 215, 525–531.

18. B. Li, S. Luan, J. Chen, Y. Zhou, T. Wang, Z. Li, Y. Fu, A. Zhai and C. Bi, Molecular Therapy-Nucleic Acids, 2020, 19, 814–826.

19. E. Potolitsyna, S. Hazell Pickering, T. Germier, P. Collas and N. Briand, Scientific reports, 2022, 12, 10157.

20. R. Li, W. Zhang, Z. Yan, W. Liu, J. Fan, Y. Feng, Z. Zeng, D. Cao, R. C. Haydon and H. H. Luu, Aging (Albany NY), 2021, 13, 4199.

21. M. Li, Z. Xie, P. Wang, J. Li, W. Liu, S. a. Tang, Z. Liu, X. Wu, Y. Wu and H. Shen, Cell death & disease, 2018, 9, 554.

22. C. Ju, R. Liu, Y.-W. Zhang, Y. Zhang, R. Zhou, J. Sun, X.-B. Lv and Z. Zhang, Biomedicine & Pharmacotherapy, 2019, 115, 108912.

23. X. Xue, X. Hong, J. Fu and C. Deng, Biophysical Journal, 2016, 110, 134a.

24. X. Xue, X. Hong, Z. Li, C. X. Deng and J. Fu, Biomaterials, 2017, 134, 22–30.

25. Y. Zhao, K. Richardson, R. Yang, Z. Bousraou, Y. K. Lee, S. Fasciano and S. Wang, Frontiers in Bioengineering and Biotechnology, 2022, 10, 1007430–1007430.

26. A. E. Adeniran-Catlett, L. D. Weinstock, F. K. Bozal, E. Beguin, A. T. Caraballo and S. K. Murthy, Biotechnology Progress, 2016, 32, 440–446.

27. J. Choi, S. Y. Lee, Y.-M. Yoo and C. H. Kim, Cell biochemistry and biophysics, 2017, 75, 87–94.

28. H.-W. Wu, C.-C. Lin, S.-M. Hwang, Y.-J. Chang and G.-B. Lee, Microfluidics and nanofluidics, 2011, 11, 545–556.

29. M. Wiewiora, J. Piecuch, M. Glűck, L. Slowinska-Lozynska and K. Sosada, Clinical hemorheology and microcirculation, 2013, 54, 313–323.

30. S. Fasciano, S. Luo and S. Wang, The Analyst, 2023, 148, 6261–6273.

31. S. Arora, A. Srinivasan, C. M. Leung and Y.-C. Toh, Current stem cell research & therapy, 2020, 15, 414–427.

32. M. Knippenberg, M. N. Helder, B. Zandieh Doulabi, C. M. Semeins, P. I. Wuisman and J. Klein-Nulend, Tissue engineering, 2005, 11, 1780–1788.

33. Y. Gao, F. Xiao, C. Wang, C. Wang, P. Cui, X. Zhang and X. Chen, Journal of Cellular Biochemistry, 2018, 119, 6986–6996.

34. J. Wu, J. Zhao, L. Sun, Y. Pan, H. Wang and W.-B. Zhang, Bone, 2018, 108, 62–70.

35. Q. Liu, X. Hu, X. Zhang, L. Dai, X. Duan, C. Zhou and Y. Ao, Molecular Therapy, 2016, 24, 1726–1733.

36. R. Riahi, Z. Dean, T.-H. Wu, M. A. Teitell, P.-Y. Chiou, D. D. Zhang and P. K. Wong, The Analyst, 2013, 138, 4777–4785.

37. R. Riahi, S. Wang, M. Long, N. Li, P.-Y. Chiou, D. D. Zhang and P. K. Wong, ACS nano, 2014, 8, 3597–3605.

38. S. Wang, R. Riahi, N. Li, D. D. Zhang and P. K. Wong, Advanced materials, 2015, 27, 6034–6038.

39. S. Tao, S. Wang, S. J. Moghaddam, A. Ooi, E. Chapman, P. K. Wong and D. D. Zhang, Cancer research, 2014, 74, 7430–7441.

40. S. Fasciano and S. Wang, SLAS technology, 2023.

41. S. Wang, S. Majumder, N. J. Emery and A. P. Liu, Synthetic Biology, 2018, 3, ysy005.

42. S. Wang, J. Sun, D. Zhang and P. Wong, Nanoscale, 2016, 8, 16894–16901.

43. S. Wang, J. Sun, Y. Xiao, Y. Lu, D. D. Zhang and P. K. Wong, Advanced Biosystems, 2017, 1.

44. Y. Zhao, R. Yang, Z. Bousraou, K. Richardson and S. Wang, Scientific reports, 2022, 12, 10315.

45. Z. S. Dean, R. Riahi and P. K. Wong, Biomaterials, 2015, 37, 156–163.

46. Y. Zhao and S. Wang, in MicroRNA Detection and Target Identification: Methods and Protocols, Springer, 2023, pp. 75–87.

47. Z. S. Dean, P. Elias, N. Jamilpour, U. Utzinger and P. K. Wong, Analytical chemistry, 2016, 88, 8902–8907.

48. G. Pachón-Peña and M. A. Bredella, Trends in Endocrinology & Metabolism, 2022, 33, 401–408.

49. M. Tencerova, M. Ferencakova and M. Kassem, Best Practice & Research Clinical Endocrinology & Metabolism, 2021, 35, 101545.

50. T. Liu, G. Melkus, T. Ramsay, A. Sheikh, O. Laneuville and G. Trudel, Nature communications, 2023, 14, 4799.

51. H. Wang, Y. Leng and Y. Gong, Frontiers in endocrinology, 2018, 9, 694.

52. F. J. de Paula and C. J. Rosen, Annual review of physiology, 2020, 82, 461–484.

53. S. Weinbaum, S. C. Cowin and Y. Zeng, Journal of biomechanics, 1994, 27, 339–360.

54. C. Price, X. Zhou, W. Li and L. Wang, Journal of Bone and Mineral Research, 2011, 26, 277–285.

55. Y. Zhao, J. Ning, H. Teng, Y. Deng, M. Sheldon, L. Shi, C. Martinez, J. Zhang, A. Tian and Y. Sun, Nature communications, 2024, 15, 2384.

56. W. Liu, Y. Zhang, Q. Li, X. Wang, Y. Wu, H. Shen and P. Wang, Pathology-Research and Practice, 2024, 155413.

57. C. Jiang, P. Wang, Z. Tan and Y. Zhang, Open Life Sciences, 2024, 19, 20220908.

58. G. Chen, C. Deng and Y.-P. Li, International journal of biological sciences, 2012, 8, 272.

59. L. Hu, W. Chen, A. Qian and Y.-P. Li, Bone Research, 2024, 12, 39.

60. J. Yang, P. Andre, L. Ye and Y.-Z. Yang, International journal of oral science, 2015, 7, 73–79.

61. M. I. Dishowitz, S. P. Terkhorn, S. A. Bostic and K. D. Hankenson, Journal of Orthopaedic Research, 2012, 30, 296–303.

62. A. Stellpflug, J. Caron, S. Fasciano, B. Wang and S. Wang, Nanoscale Advances, 2025.

