## Supplementary material for "Shear stress inhibits adipocyte differentiation via downregulating lncRNA MALAT1": Fig. S1-S5

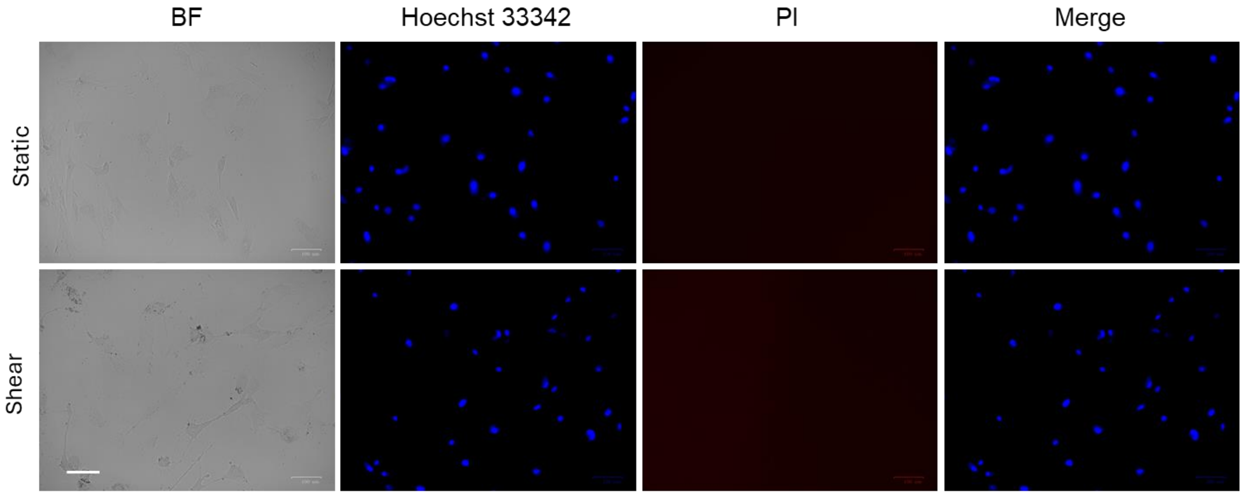


**Figure S1.** Representative images of MSCs under static and shear conditions after 5 days of culture. BF: Bright field, Hoechst 33342: Nucleus staining. Propidium Iodide (PI): stain dead cells. (n=3) Scale bar: 100 μm.

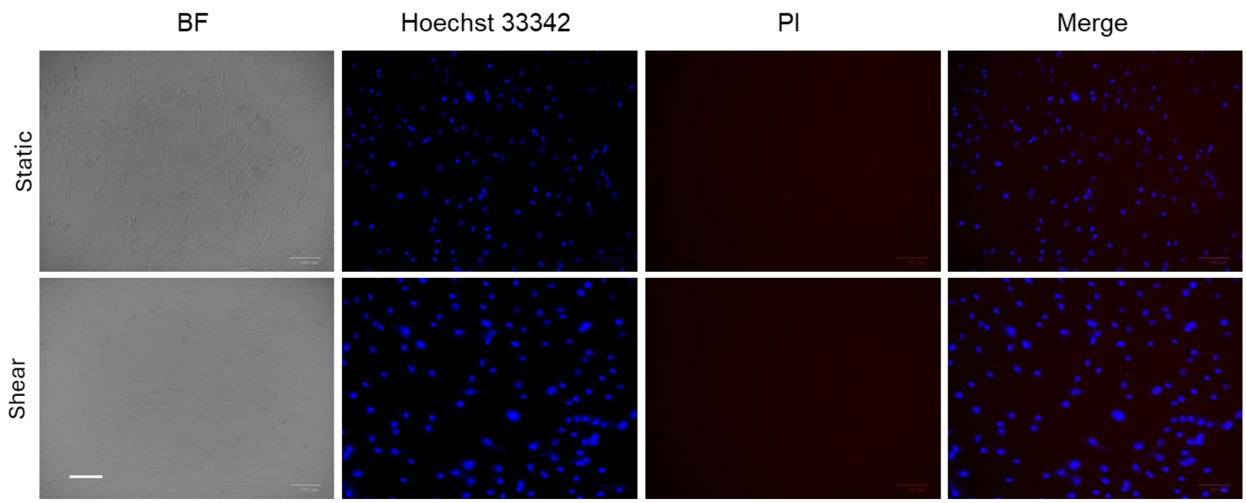


**Figure S2.** Representative images of MSCs under static and shear conditions after 10 days of culture. BF: Bright field, Hoechst 33342: Nucleus staining. Propidium Iodide (PI): stain dead cells. (n=3) Scale bar: 100 μm.

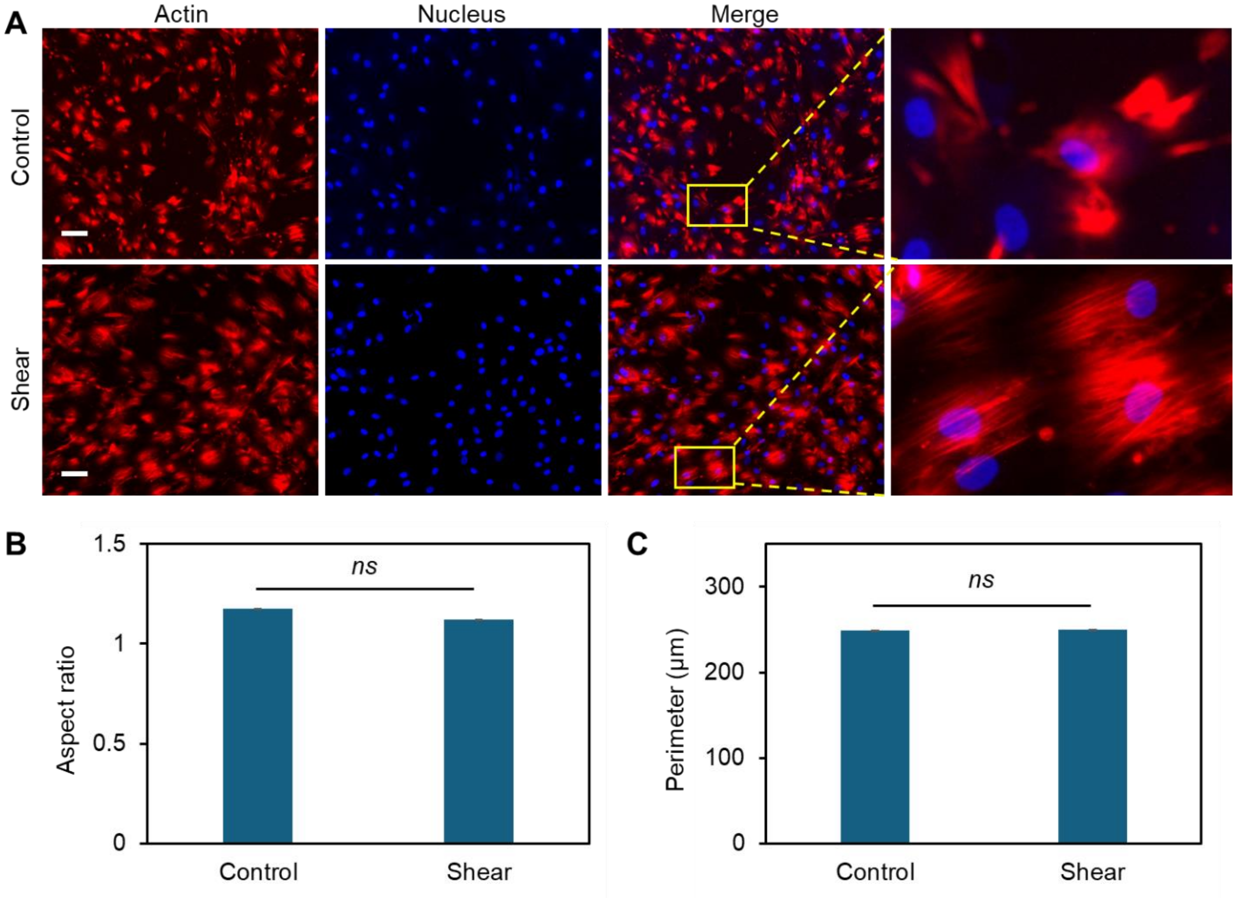


**Figure S3. Effects of shear stress on MSCs morphological changes and actin organization**. **(A)** Representative bright field and fluorescence images of MSCs under static condition (control) and exposed to shear stress (shear). MSCs were exposed to orbital shear continuously for 5 days. Samples were stained with F-actin (red; by phalloidin), and nuclei (blue; by Hoechst 33342), respectively. Scale bar: 100 µm. Comparison of aspect ratio **(B)** and perimeters **(C)** of MSCs after 10 days of exposure to low fluid shear. Data represents over 200 cells in each group and are expressed as mean± s.e.m. (n = 5, ***, $p < 0.001$, **, $p < 0.01$, *, $p < 0.05$).

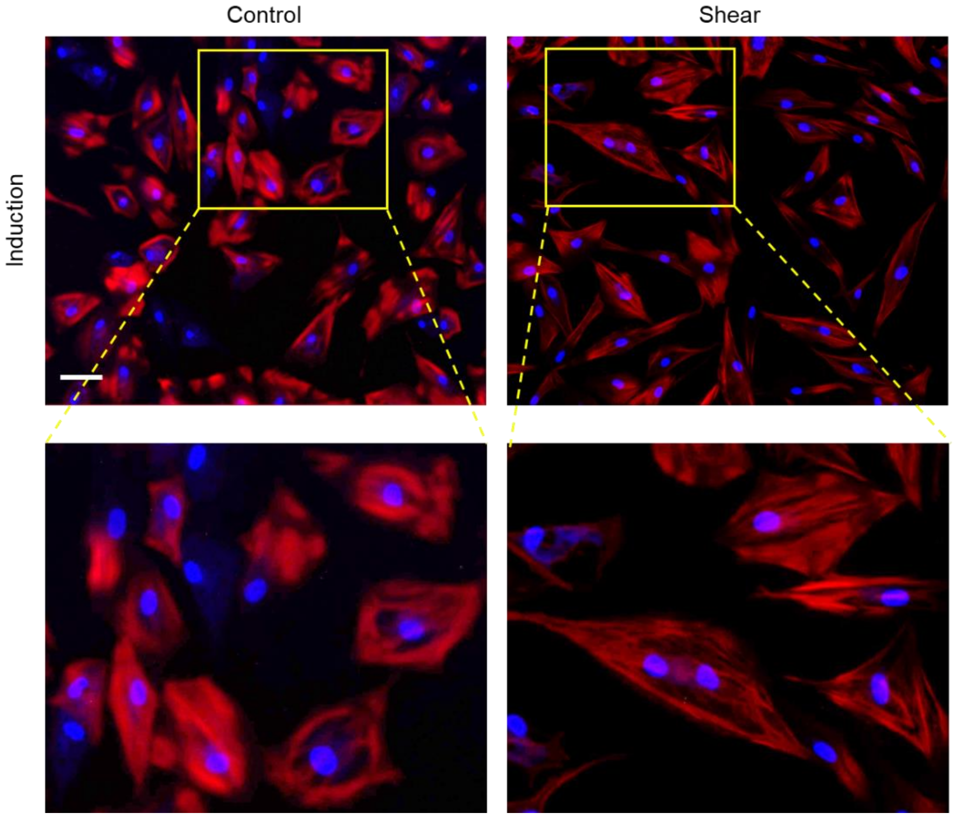


**Figure S4. Effects of shear stress on actin organization during adipocyte differentiation**. Red: F-actin. Blue: Nucleus. During adipocyte differentiation, actin cytoskeleton under drastic remodeling from aligned stress fibers to cortical actin structures. For cells exposed to shear stress, actin cytoskeleton remodeling was mediated and stress fibers maintained elongated inside cells.

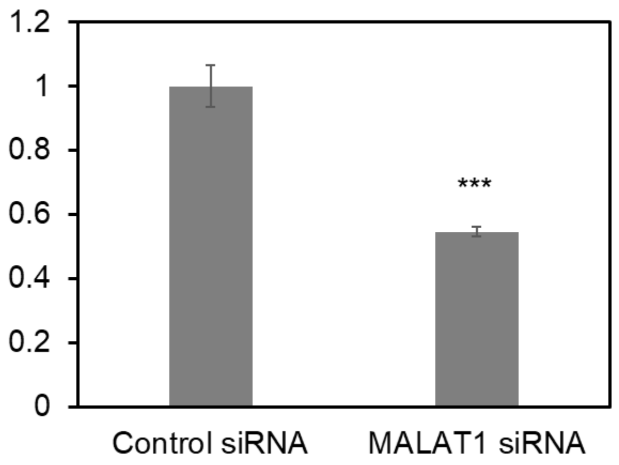


**Figure S5. MALAT1 siRNA knockdown efficiency.** RT-PCR was performed to evaluate the silencing efficiency. The relative expression levels of MALAT1 were determined by the equation $2^{-\Delta\Delta Ct}$. Data are expressed as mean ± s.e.m. (n = 3). A two-tailed t-test was used to analyze differences between control siRNA and MALAT1 siRNA. ***, $p < 0.005$.

**Table S1. MALAT1 LNA probe sequences**

| Name | | Sequence (5'-3') | Fluorophore |
|---|---|---|---|
| MALAT1 | Donor | +T+C+G+C+A TACGT GTGTC TGCTG AGTGT +T+C+C+T+G | /56-FAM |
| | Quencher | +G+A+C+A+C ACGTA TGCGA | /3-Iowa BlackFQ |
